# Degron-modified Cas12a enhances single-cell CRISPR screening

**DOI:** 10.1101/2024.12.08.627374

**Authors:** Valentina Snetkova, Carolina Galan, Romain Lopez, Antonio R. Rios, Takamasa Kudo, Kristel Dorighi, Søren Warming, Benjamin J. Haley

## Abstract

Single-cell CRISPR (Perturb-seq) screens have primarily relied on Cas9 whereas Cas12a, despite its unique effectiveness for multiplex guide expression, remains underexplored. This may be due to Cas12a’s guide RNA array (pre-crRNA) self-processing activity and the subsequent challenges associated with pre-crRNA sequence recovery. By developing modified pre-crRNA expression vectors and a degron-based Cas12a system, we overcome the self-processing constraint, allowing for accurate detection of pre-crRNAs at the single-cell level, thus greatly expanding possibilities for future Perturb-seq efforts.

## Main

The rapidly evolving CRISPR toolbox has transformed functional genomics by offering efficient, versatile, and complementary approaches for high-throughput genetic modification^1–4^. Efforts have been made to broaden the application of functional genomic screens through targeting of multiple genes in each cell and/or to simplify the experimental process by reducing the number of constructs necessary for a screening library^5–7^. CRISPR-associated protein 12a (Cas12a, previously Cpf1) has emerged as a useful nuclease for these purposes^8–11^. As with Cas9 and other Class II CRISPR enzymes, Cas12a can be programmed with a short guide RNA (crRNA) to direct site-specific DNA breaks. However, unlike Cas9, it has a native ability to process an expressed guide RNA array (pre-crRNA)^12^. This minimizes the burden of cloning plasmid libraries intended to express several independent Cas12a crRNA sequences from each vector, which can then disrupt one or many genes at a time^13,14^. Engineered variants of Cas12a that boost specificity and activity have further enhanced its capabilities as a gene perturbation technology^15,16^.

The power of functional genomic screens has dramatically increased through direct integration of CRISPR-mediated gene perturbation and single-cell (sc) assay technologies (*e.g.* scRNA-seq), an approach which has since been broadly termed Perturb-seq^17–21^. Upon delivery of guide RNAs into a cell population, expression changes for thousands of mRNAs in response to numerous individual gene-targeting events can be measured with single cell resolution. This enables detection of rich and heterogeneous cellular phenotypes that go beyond individual pathway reporters or overt viability effects used for standard, pooled vector-based screens.

Perturb-seq has primarily relied on Cas9 for gene targeting. To harness the unique multi-guide expression capabilities of Cas12a for single-cell applications, we developed a comprehensive suite of pre-crRNA and enhanced *Acidaminococcus sp*. (enAs)Cas12a expression vectors, along with refined methods for sequencing library preparation and analysis. We demonstrate effective targeting of at least four genes from a single vector and identify transcriptomic signatures that result from disruption of select epigenetic factors using a custom, minimal pre-crRNA library. Together, this new platform will expand the scalability and complexity of future Perturb-seq screening campaigns.

To adapt Cas12a for Perturb-seq, we generated a population of NCI-H358 (H358, bronchioalveolar carcinoma) cells to stably express a 4XNLS-enAsCas12a transgene. We then explored adaptations of the direct-capture^21^ and CROP-seq^19^ strategies for lentiviral delivery and subsequent single cell detection of pre-crRNAs (Extended Data Fig. 1a). For Cas9-based direct-capture, one of two different capture sequences (CS 1 or 2) is incorporated within a stem loop of the single-guide (sg)RNA scaffold or is appended to the sgRNA 3’ terminus^21^. Due to differences between the Cas9-associated sgRNAs and the relatively shorter, reversed orientation of Cas12a-associated crRNAs, we tested three different pre-crRNA configurations with varying CS positions: at the 3’ end of the spacer (position 1), between the direct repeat (DR) and the spacer (position 2), and in the hairpin of the DR (position 3) (Extended Data Fig. 1b). The effectiveness of each approach was tested through expression of a representative pre-crRNA array designed to simultaneously disrupt either the CD63 and CD47 or CD81 and CD9 genes (one crRNA per target, two targets per vector), respectively, followed by assessment of cell surface levels for each gene product (Extended Data Fig. 1c, d). The pre-crRNA engineered with a CS in position 1 showed comparable activity to an unmodified pre-crRNA, while CSs in the two other positions interfered with Cas12a targeting (Extended Data Fig. 1e). Modest, context-dependent differences on targeting activity were observed with constructs incorporating CS1 vs. CS2, suggesting capture sequence or experiment-specific effects could impact function.

Having established the optimal CS position, we compared the efficacy of pre-crRNAs expressed from the direct-capture format vs. otherwise unmodified pre-crRNAs within a CROP-seq vector. In the CROP-seq context, the pre-crRNA is adjacent to the lentiviral 3’ long terminal repeat (LTR) and an RNA Pol-II termination sequence (Extended Data Fig 2a). This enables the guides to be functionally expressed from an RNA Pol-III (U6) promoter formed from a duplication event during lentiviral integration. In addition, the pre-cRNA will be incorporated into an RNA Pol-II (*EF1A,* or *EEF1A1*)-derived mRNA necessary for antibiotic (puromycin) selection marker expression. This mRNA is later amplified and the pre-crRNAs identified following oligo(d)T priming and 3’ scRNA-seq^19,22^. The pre-crRNA is positioned away from the 3’ LTR in the direct capture context, which excludes the ability to identify guides using conventional 3’ or 5’ (*i.e.* SMART) scRNA-seq protocols^23^. For relative assessment, we designed pre-crRNAs to be expressed from either the direct-capture Perturb-seq or CROP-seq formats for simultaneous, multiplexed disruption of either two (CD47/CD63 or CD81/CD9) or four genes (CD47/CD63/CD81/CD9) (Extended Data Fig. 2b). Comparable perturbation of multiple targets resulted from each of the pre-crRNA vector contexts (Extended Data Fig. 2c). There was no overt difference in activity levels when individual crRNAs were reordered within the array, although efficacy was impacted by the spacer sequence for a given target gene (Extended Data Fig. 2c-d, and see Supplemental Note 1).

Epigenetic modifiers have recognized roles in controlling gene expression and influencing cell fate changes, which can be monitored by scRNA-seq^24^. Accordingly, we selected a subset of these factors for assessment with pooled enAsCas12a CROP-seq or direct-capture libraries (Fig. 1a). For both formats, the vector library included identical crRNAs targeting 45 total factors that were reported to be highly-expressed in H358 cells^25^ and play diverse roles such as chromatin remodeling (*e.g.*, ACTL6A, CHD3, SMARCA4), histone modification (*e.g.*, EHMT1, EHMT2, KAT6A, KDM1A), and DNA methylation (*e.g.*, DNMT1, MECP2) (Fig. 1b). A single vector was generated per gene, with each gene targeted by a pair of crRNAs. Based on prior experience with single vs. multi-guide targeting using Cas9^26^, we reasoned that two unique crRNAs directed towards the same gene would increase the KO efficiency when co-expressed with enAsCas12a.

**Fig. 1.**
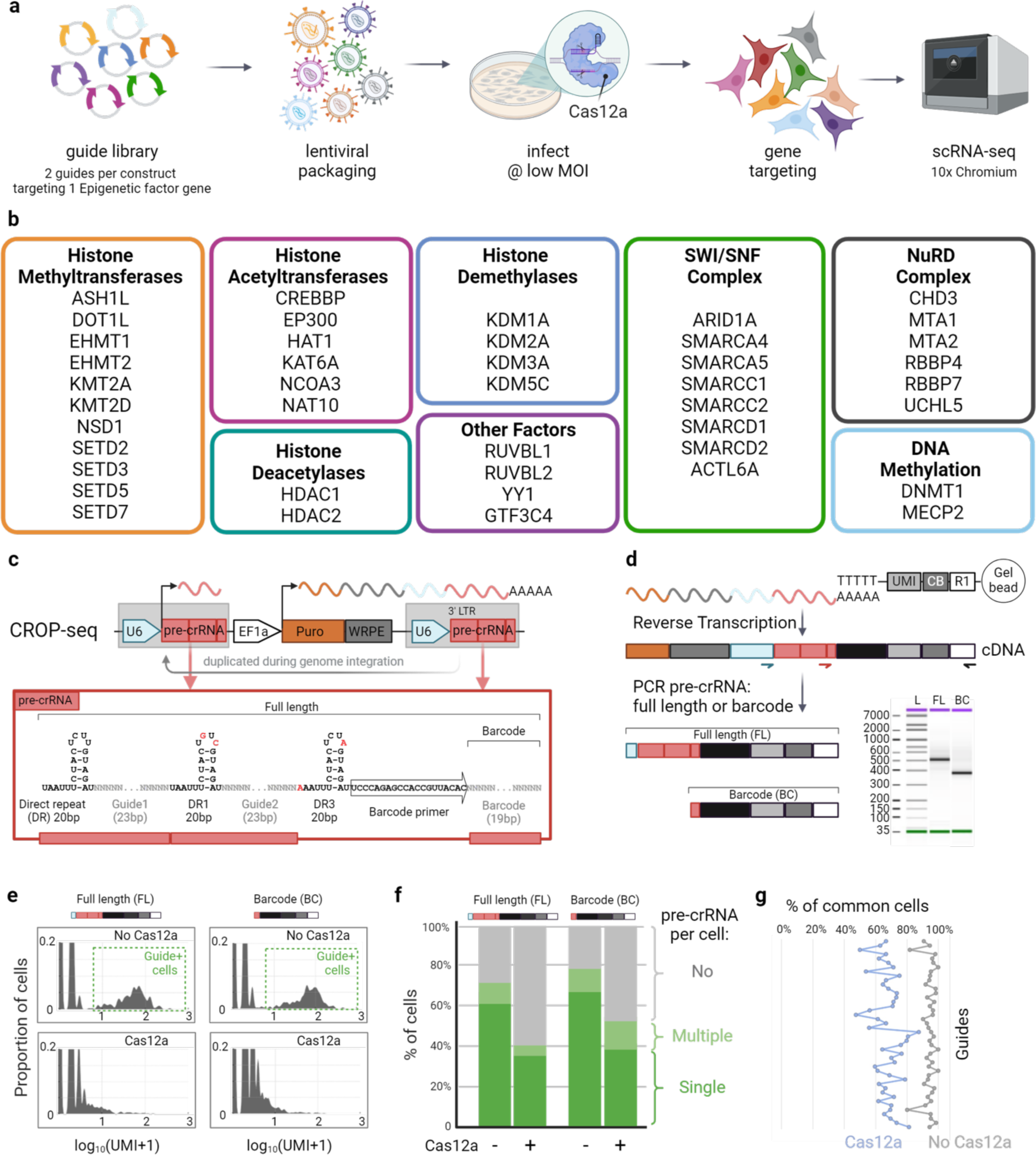
Cas12a impairs guide detection in scRNA-seq. **a,** Experimental strategy for Cas12a CRISPR screen with scRNA-seq read-out. **b,** Epigenetic factors selected for the screen. Factors were selected by gene expression in H358 cells and diversity of function. Libraries contained pre-crRNAs with two guides for each factor. **c,** pre-crRNA construct in CROP-seq format. **d,** Protocol for amplifying pre-crRNA sequences from single-cell cDNA showing representative analytical gel electrophoresis. **e,** Signal from representative pre-crRNA obtained by full-length and barcode detection methods. **f,** Percentage of cells identified as expressing none, single, or multiple pre-crRNA constructs. **g,** Percentage of cells with consistent pre-crRNA identify calls between full-length and barcode methods.

Beginning with the CROP-seq format library, we appended a 3’ 19bp barcode separated by a direct repeat and a barcode segment-specific primer site to each pre-crRNA array to assist with both pre-crRNA capture and accurate single cell assignment of each crRNA pair (Fig. 1c, d). Sequencing amplicon preparation was optimized for either the full-length or barcode-only regions of the pre-crRNA cassettes (Fig. 1d). To map the full-length pre-crRNA sequences, we developed a computational method based on creation of a custom pre-crRNA transcriptome with the Cell Ranger algorithm^27^, whereas mapping and assignment of cells to each barcode was performed using a standard scRNA-seq analysis pipeline (Extended Data Fig. 3).

Next, we compared recovery of pre-crRNAs from cells transduced with the CROP-seq library in the presence or absence of enAsCas12a. Library transduction conditions were optimized for integration of ∼1 vector per cellular genome, and this was followed by four days in culture after antibiotic selection to facilitate pre-crRNA expression and target gene disruption. The barcode sequencing strategy enabled relatively higher fraction assignment of pre-crRNAs to individual cells when compared to the full-length pre-crRNA approach (Fig. 1e,f). However, in cell populations expressing enAsCas12a, we detected substantially fewer pre-crRNAs and reduced consistency in assignment of pre-crRNAs within each cell, regardless of the method for sequencing library preparation (Fig. 1e-g). To account for potential vector-specific influences on pre-crRNA recovery, we performed a similar series of screens and sequencing studies with an otherwise equivalent direct-capture library. This format showed inadequate pre-crRNA recovery and was deprioritized for further assessment (Extended Data Fig. 4a, b and see Supplementary Note 2). We hypothesized that Cas12a’s self-processing activity was leading to degradation of pre-crRNA-containing transcripts and, as a result, poor assignment of perturbation sequences within individual cells.

Elimination of enAsCas12a from library-transduced cells following gene disruption could provide an opportunity for pre-crRNA accumulation and improved recovery in single cells. As a simple, pharmacological means to accomplish this, we leveraged the degradation tag (dTag) system^28^. Within a standard lentiviral vector, an enAsCas12a transgene was engineered to be expressed with a C-terminal FKBP12^F36V^ fusion domain (enAsCas12a-FKBP12^F36V^) to enable selective ubiquitination and, subsequently, degradation upon addition of a dTag small molecule (*e.g.* dTagV1). After validating drug-induced degradation of the enAsCas12a-FKBP12^F36V^ protein in stably-expressing H358 cells (Extended Data Fig. 5), we tested the impact of this fusion on gene targeting efficiency in the presence or absence of dTagV1. As before, lentivirus-delivered pre-crRNAs were used to deplete CD81 and CD9 or CD47 and CD63, this time in cells expressing either enAsCas12a or enAsCas12a-FKBP12^F36V^ (Fig 2a). In the absence of dTagV1, enAsCas12a and enAsCas12a-FKBP12^F36V^ showed similar levels of targeting efficiency (Fig 2b). Continuous culture of enAsCas12a-FKBP12^F36V^ with dTagV1 (before and after pre-crRNA delivery) prevented knock-out for three out of four genes; only CD9 receptor levels were reduced in cells treated with dTagV1. This may be due to residual but undetectable enAsCas12a combined with a highly-potent *CD9*-targeting crRNA and/or potential differences in turnover or membrane trafficking kinetics of CD9 protein relative to the other targets we evaluated.

**Fig. 2.**
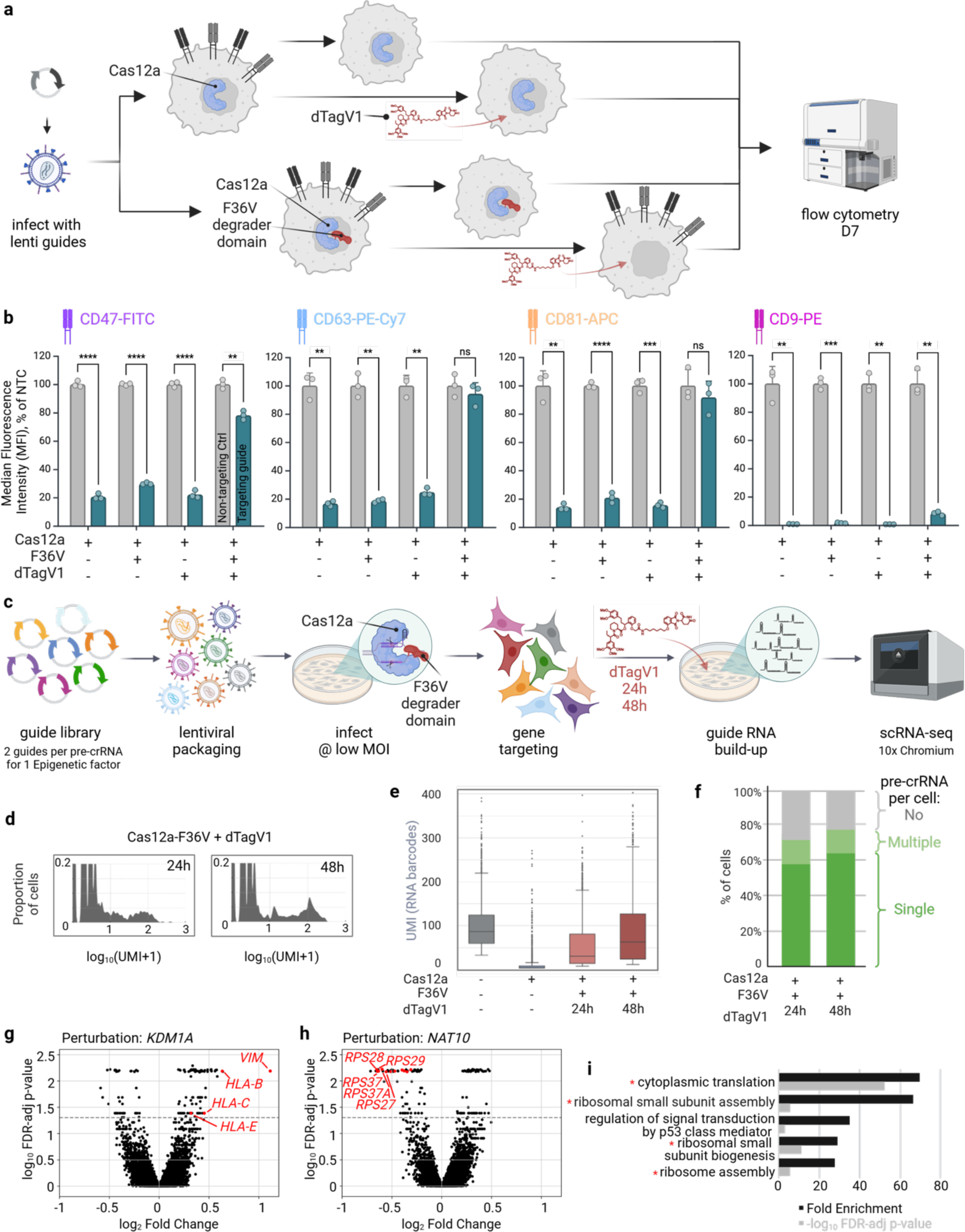
Degradation of Cas12a restores pre-crRNA detection. **a**, Experimental strategy. **b**, Quantification of FACS data comparing Cas12a-FKBP12^F36V^ and Cas12a-mediated editing efficiency and prevention of editing in the presence of dTagv1. Bar heights represent the Mean of three biological replicates normalized by average NTC values. Error bars represent SD. Statistical comparison was performed by an unpaired two-sample t-test with Welch’s correction whereby *p < 0.05, **p < 0.01, ***p < 0.001, ****p < 0.0001 as compared to the NTC. **c**, Experimental strategy for scRNA-seq with Cas12a-FKBP12^F36V^ and incorporation of dTagV1 addition to build-up pre-crRNA post gene targeting. **d**, UMI signal from representative pre-crRNA obtained from barcodes after 24h or 48h dTagV1 treatment. **e**, Boxplot of UMI signals for all pre-crRNA in the library obtained from barcodes after 24h or 48h dTagV1 treatment. Statistical comparison between samples was performed by an Mann-Whitney U test whereby the p-values were 0 (Cas12a), 3.07e^-279^ (24h), and 1.32e^-61^ (48h), as compared to the first sample (no Cas12a). **f**, Percentage of cells identified as expressing none, a single or multiple pre-crRNAs based on the barcode sequence from the epigenetic factor screen described in c. **g**, **h**, Volcano plots of transcriptome-wide changes in cells with *KDM1A or NAT10*-targeting pre-cRNA vs NTCs. Genes highlighted in red have been reported in the literature. **i**, Top 5 biological pathways from gene ontology analysis of significantly altered genes shown in h with red asterisks highlighting the pathways reported to be affected by *NAT10* perturbation before. Black bars represent fold enrichment values, while grey bars show -log_10_ FDR-adjusted p-values.

After establishing that enAsCas12a-FKBP12^F36V^ maintained high gene editing capacity, we repeated the epigenetic factor screen with dTagV1-mediated depletion of enAsCas12a-FKBP12^F36V^ for 24 or 48 hours before collecting samples for single cell sequencing (Fig. 2c). Analysis of pre-crRNA counts showed continued improvement in guide recovery over the time points tested with dTagV1 treatment (Fig. 2d, e). For both the 24- and 48-hour treatments we were able to recover and assign pre-crRNAs to individual cells with frequencies similar to library-infected populations devoid of enAsCas12a (up to 80%) (Fig. 1f, 2f). In line with a generalized negative influence from enAsCas12a expression on pre-cRNA recovery, we observed similar results with full-length pre-crRNA counts (Extended Data Fig. 6a, b, c) in addition to improved consistency when matching pre-crRNAs to individual cells with either the full-length or barcode only methods (Extended Data Fig. 6d). Increased pre-crRNA recovery was observed following a 48-hour pulse with dTagV1 in a separate cell line (K562, chronic myeloid leukemia) engineered to express CROP-seq-formatted pre-crRNAs and enAsCas12a-FKBP12^F36V^ (Extended Data Fig. 7). These results demonstrate the utility of decreasing enAsCas12a levels immediately prior to Perturb-seq sample collection.

Expression profiles following disruption of lysine demethylase 1A (KDM1A/LSD1) and N-acetyltransferase 10 (NAT10) exemplify both the specificity and efficacy of phenotypes generated using the dual crRNA library (Fig 2g, h); a heatmap summarizing gene expression changes for all perturbations can be found in Extended Data Fig 8. KDM1A is known to directly inhibit expression of vimentin (VIM)^29,30^, which represents the most upregulated mRNA in cells with the *KDM1A* pre-crRNA (Fig. 2g). We also observed upregulation of individual genes within the HLA-I family (*i.e. HLA-B, C*, and *E*) in the same cells (Fig. 2g). It was recently shown that pharmacological targeting of KDM1A induced strong expression of several HLA-I genes in a panel of Small Cell Lung Cancer lines, with the increase in *HLA-I* expression occurring only after seven or more days of culture in the presence of the inhibitor^31^. While this coincides with the time point selected for our screen, it is possible that these (and other differences in gene expression observed in our screen) are secondary effects of target disruption, and distinct changes might be detected through an earlier time course analysis. Separately, consistent with its role in promoting ribosome biogenesis^32^, 15 of the top 25 most down-regulated mRNAs in cells containing the *NAT10* pre-crRNA array encode for unique ribosomal subunits (Fig. 2h), which was also reflected in gene ontology analysis where 4 out of top 5 biological pathways were related to translation (Fig. 2i).

By establishing an enhanced, multiplexed pre-crRNA expression cassette and chemically-degradable enAsCas12a variant, we overcome existing barriers to single cell guide RNA recovery and assignment for functional genomic screens that make use of this uniquely-capable DNA nuclease. In addition, we developed optimized methods for library preparation enabling either full-length pre-crRNA sequencing or guide RNA identification via a pre-crRNA-linked barcode. A proof-of-concept, loss-of-function screen targeting a subset of epigenetic factors demonstrated that our pre-crRNA expression vector library and degradable enAsCas12a system facilitated precise and efficient gene disruption. While our approach employed the FKBP12^F36V^ domain and a corresponding dTag molecule for induced degradation of enAsCas12a, emerging degron technologies, including direct targeting with a PROTAC molecule, could offer greater flexibility and more effective Cas12a reduction before sample collection for sequencing^33,34^. Overall, our framework for the application of enAsCas12a-mediated Perturb-seq significantly amplifies the potential and broadens the opportunity space for this powerful genetic screening technology.

## Methods

### Cell culture

H358 cells (ATCC CRL-5807^™^) and K562 (ATCC CCL-243^™^) were sourced from Genentech’s internal cell bank, where the cell lines were maintained under mycoplasma-free conditions. Both cell lines were cultured in RPMI-1640 growth medium + 10% heat inactivated FBS (Sigma, F4135) + 2mM L-Glutamine and 100 U/mL Penicillin-streptomycin (Gibco, 15140-122).

### Lentiviral production and cell transduction

Plasmids expressing desired cargo (enAsCas12a or pre-crRNA) and lentiviral packaging constructs (VSVg and Delta8.9) were transiently co-transfected into HEK293T cells with Lipofectamine 2000. After 48h, supernatants were collected, filtered through a 0.45µM PES filter and concentrated with Lenti-X Concentrator (Takara, 631232) according to the manufacturer’s manual.

Transduction was performed in 8 µg/ml polybrene (Sigma Cat#TR-1003r) with direct addition of filtered and concentrated viral supernatant. Transduced cells were selected with 15 µg/ml blasticidin (for enAsCas12a constructs) or 1 µg/ml puromycin (for pre-crRNA constructs and libraries). MOI of lentiviral infection was assessed with CellTiter-Glo Luminescent Cell Viability Assay (Promega, G7572).

### Vector Sequences

Transgene sequences for enAsCas12a, enAsCas12a-FKBP12^F36V^ and pre-crRNA sequences are detailed in the Supplementary Tables. crRNA sequences for CD47 and CD63 were sourced from the literature^9^. crRNA sequences for CD81, CD9, epigenetic factors, and non-targeting controls (NTCs) were designed using crisprVerse Bioconductor ecosystem^35^ using the GRCh38 genome build. Sequences not found in the human genome or targeting olfactory receptors were used as non-targeting controls (NTCs) to account for non-cutting and cutting but non-targeting activity. Different direct repeats were used in the pre-crRNA containing multiple crRNAs to limit possible recombination^9^. pre-crRNA sequences were produced as DNA oligonucleotides and cloned into a puromycin-selectable lentiviral vector pLKO_SHC201 (Sigma) by Genscript. For libraries, final plasmids were pooled together in equimolar ratios, concentrated (Millipore, UFC5010), and column purified (Qiagen, 28004).

### Flow cytometry

For surface protein analysis, staining was performed using the following antibodies: anti-human CD47 (FITC, Cat No. 323106), anti-human CD63 (PE-Cy7, Cat No. 353010), anti-human CD81 (APC, Cat No. 17-0819-42), and anti-human CD9 (PE, Cat No. 312106). Each antibody was used at a 1:25 dilution. The staining was conducted for ∼60 minutes on ice in the MACS buffer. Then, cells were washed twice with the MACS buffer and fixed with 1% PFA for 5 minutes at RT to inactivate any remaining lentiviral particles, followed by a final wash with the MACS buffer. Flow cytometry was performed on a BD FACSymphony instrument and data was analyzed using FlowJo software, plotted in BioRender and statistical analysis performed with R 4.0.0.

### Single-cell RNA sequencing

Plasmid libraries consisting of 50 dual pre-crRNAs each (45 with two crRNA on one pre-crRNA targeting the same epigenetic factor and 5 NTCs) were transduced into H358-enAsCas12a or H358-enAsCas12a-FKP12^F36V^ cells at targeted MOI of 0.3 and a minimum coverage of 1000x cells per pre-crRNA. 1ng/µl puromycin selection was initiated 48h post-transduction. Some cells were split before puromycin selection to confirm MOI with Cell-Titer-Glo (Promega, G7570). The confirmed MOI was 0.27 for the CROP-seq library and 0.38 for the direct-capture Perturb-seq library in the H358-enAsCas12a cells; and 0.29 for the CROP-seq library in H358-enAsCas12a-FKBP12^F36V^ cells.

### Cell processing and library preparation

In the case of H358-enAsCas12a cells, after the introduction of the targeting library and initiation of the puromycin, cells were cultured for a week, then dissociated, washed in 1xPBS, counted to ensure high viability (>90%) and resuspended in 1% BSA in 1xPBS to process through the 10x Chromium Controller according to the manufacturer’s protocol (CG000316 or CG000418) with a recovery aim of 10,000 - 20,000 cells per lane. In the case of H358-enAsCas12a-FKBP12^F36V^, prior to 10x processing, cells were cultured in 1µM dTAGV-1 (Tocris Bioscience, 6914) for 24h or 48h. dTagV1 was continuously supplied until the step of reverse transcription. Gene expression libraries were prepared according to the 10x protocol. In the cases when samples were hashed, the cells were labeled with the TotalSeq^TM^ anti-human Hashtag antibodies from BioLegend (HTO #1-4,6-9, cat. # 394661, 394603, 394605, 394637, 394641, 394673, 394675, 394677) according to the manufacturer’s instructions. The hashtag (HTO) libraries were amplified with addition of a primer 5’-GTGACTGGAGTTCAGACGTGTGCTCTTCCGAT*C*T at 2nM final concentration to the cDNA amplification step in the 10x protocol. Subsequently HTO libraries were amplified from lower molecular weight portion (pellet) of amplified cDNA with Dual Index TT Set A (10x Genomics, PN-1000215) in the conditions mimicking indexing of GEX libraries.

### Pre-crRNA library preparation

The CROP-seq and polyadenylated sequences from direct-capture CROP-seq constructs were amplified from cDNA with forward primer 5’-GTGACTGGAGTTCAGACGTGTGCTCTTCCGATCTtggaaaggacgaggtacc (for full-length pre-crRNA) OR 5’-GTGACTGGAGTTCAGACGTGTGCTCTTCCGATCTTCCCAGAGCCACCGTTACAC (for barcodes) and reverse primer 5’-CTACACGACGCTCTTCCGATCT with the following conditions: 250ng cDNA, 1µM final primer concentration in a 50µl reaction with NEBNext® Q5 Hot Start HiFi PCR Master Mix (NEB, M0543L) cycling at 98°C – 3’, (98°C – 30’’, 66°C – 10’’, 72°C – 30’’) x11, 72°C – 1’. 10µl from the finished PCR were used to add multiplexing indexes with 10ul of pre-mixed primers from Dual Index TT Set A (10x Genomics, PN-1000215) in a 50µl reaction with NEBNext® Q5 Hot Start HiFi PCR Master Mix (NEB, M0543L) cycling at 98°C – 3’, (98°C – 30’’, 68°C – 10’’, 72°C – 30’’) x9, 72°C – 1’. The long direct-capture CROP-seq transcripts bound to beads through capture sequences were amplified from cDNA with forward primer 5’-GTGACTGGAGTTCAGACGTGTGCTCTTCCGATCTtggaaaggacgaggtacc (for full-length pre-crRNA) and reverse primer 5’-GCAGCGTCAGATGTGTATAAGAGACAG with the same conditions as the polyadenylated transcripts amplification, except for the indexing primers, which were taken from Dual Index NT Set A (10x Genomics, PN-1000242). CRISPR libraries for direct-capture Perturb-seq and direct-capture CROP-seq constructs were created according to 10x genomics user manuals (CG000316 or CG000418).

### Sequencing

Gene expression and pre-crRNA libraries were multiplexed and sequenced on NextSeq 2000. The intended read coverage for gene expression libraries was 20,000-30,000 reads per cells, approximately 10% of that for the hashtag (HTO) libraries and a quarter of that for the pre-crRNA libraries. The gene expression libraries. HTO libraries and pre-crRNA barcode libraries were sequenced with P3 kit at R1: 28bp, R2: 90bp, and 10bp each for i7 and i5. The full-length pre-crRNA libraries were sequenced with P1 or P2 kits at varying length of R2 to cover the full-length of the pre-crRNA.

## Data analysis

### Read alignment, demultiplexing and data pre-processing

Data analysis workflow is described in Extended Data Fig. 4 Briefly, scRNA-seq reads were mapped to the reference human genome GRCh38-2020-A or a custom reference containing full length pre-crRNA sequences using Cell Ranger v7.1.0. To call 19bp barcodes from the reads we used kallisto-bustools^36^. HTO barcodes were processed using Cell Ranger v7.1.0. Demultiplexing of HTO and pre-crRNA feature barcoding data and assigning identity to cells was performed with a custom script as described in Xie *et al.*, 2019^37^. This analysis was performed for both full length pre-crRNA sequences or barcodes alone. Downstream analysis was conducted using the assigned gene expression and pre-crRNA or barcode matrix. Cells with either more than 10% mitochondrial reads or less than 3,000 UMIs were removed. Cells assigned to a perturbation with less than 15 total assigned cells were removed. Genes with less than 500 UMIs across all cells were removed, leaving a total of 12,522 genes. Single cell expression matrix and feature barcodes were processed in an anndata object format in Scanpy. Expression counts per cell were normalized to a total of 10,000 counts per cell, and normalized values were log transformed (natural log), after adding a pseudo-count of 1. Separately, Cumulus^38^ was used to create barcode and HTO matrices for data from K562 cells.

### Statistical Analysis of transcriptome-wide changes in Epigenetic factor screen data

A linear model (ordinary least-square regression) was fit for each of the 12,522 (response) genes, where the gene expression values were the response variable and the perturbation (using the NTC as a reference) as well as the total number of UMIs were used as covariates. We tested for statistical significance of the coefficient linking the perturbation identifier to the response gene expression. The false discovery rate was controlled using the Benjamini-Hochberg procedure across all the tested genes.

### Western Blot

enAsCas12a and enAsCas12a-FKP12^F36V^ -expressing NCI-H358 (H358) cells were treated with 1 µM dTAGV-1 (Tocris Bioscience, 6914) for various time points (48 hours, 24 hours, 6 hours, 2 hours, 30 minutes, and 10 minutes) or with no dTAGV-1 compound as a control (DMSO alone). Following treatment, cells were collected and lysed using RIPA Lysis and Extraction Buffer (Thermo Fisher, 89900) supplemented with cOmplete, EDTA-free Protease Inhibitor (Roche, 11873580001) according to the manufacturer’s instructions. Protein samples were prepared with 4x Laemmli Sample Buffer (Bio-Rad, 16107472) containing 2-Mercaptoethanol (Sigma, 63689) as per the manufacturer’s instructions. The samples were then separated on Novex 4-12% Tris-glycine gels (Invitrogen, XP04122BOX) using Novex™ Tris-Glycine SDS Running Buffer (10X) (Thermo Fisher, LC2675) wth Chameleon® Duo Pre-stained Protein Ladder (Licor, 928-60000) as the marker. Gels were transferred to PVDF membranes (Invitrolon™ PVDF/Filter Paper Sandwiches, 0.45 μm, 8.3 x 7.3 cm, LC2005) using Tris-Glycine Transfer Buffer (25X) (Thermo Fisher, LC3675). The membranes were then blocked with Intercept® (PBS) Blocking Buffer (Licor, 927-70001) for 2 hours at room temperature. Primary antibodies were incubated overnight at 4°C: Cas12a antibody [3D3-F7] (abcam, ab273438) at a 1:1000 dilution and β-Actin Mouse Monoclonal Antibody for normalization (Licor, 926-42212) at a 1:2000 dilution. Following overnight primary antibody incubation, membranes were washed with PBSt and incubated with IRDye® 680RD Donkey anti-Mouse IgG Secondary Antibody (Licor, 926-68072) at a 1:5000 dilution for 1 hour at room temperature. Membranes were again washed in PBS and protein bands were visualized using the Bio-Rad ChemiDoc MP Imaging System.

## Supporting information

FACS data

precrRNA sequences

## Acknowledgements

We would like to thank Dr. Shiqi Xie for his invaluable help in the initial development of the script to assign guide-cell identity in scRNA-seq and Ms. Rajini Srinivasan for help with designing the Cas12a-FKBP12^F36V^ construct.

## Author contributions

V.S. and B.J.H. conceptualized the project and designed experiments. V.S. and K.D. designed plasmids. V.S., C.G., and A.R.R. (under T.K. supervision) conducted molecular biological experiments. V.S. analyzed all the data with R.L analyzing DEG for the epigenetic factor screen. V.S., C.G., S.W., B.J.H. interpreted the data and wrote the manuscript, and all authors edited the paper.

## Extended data

**Extended Data Fig. 1.**
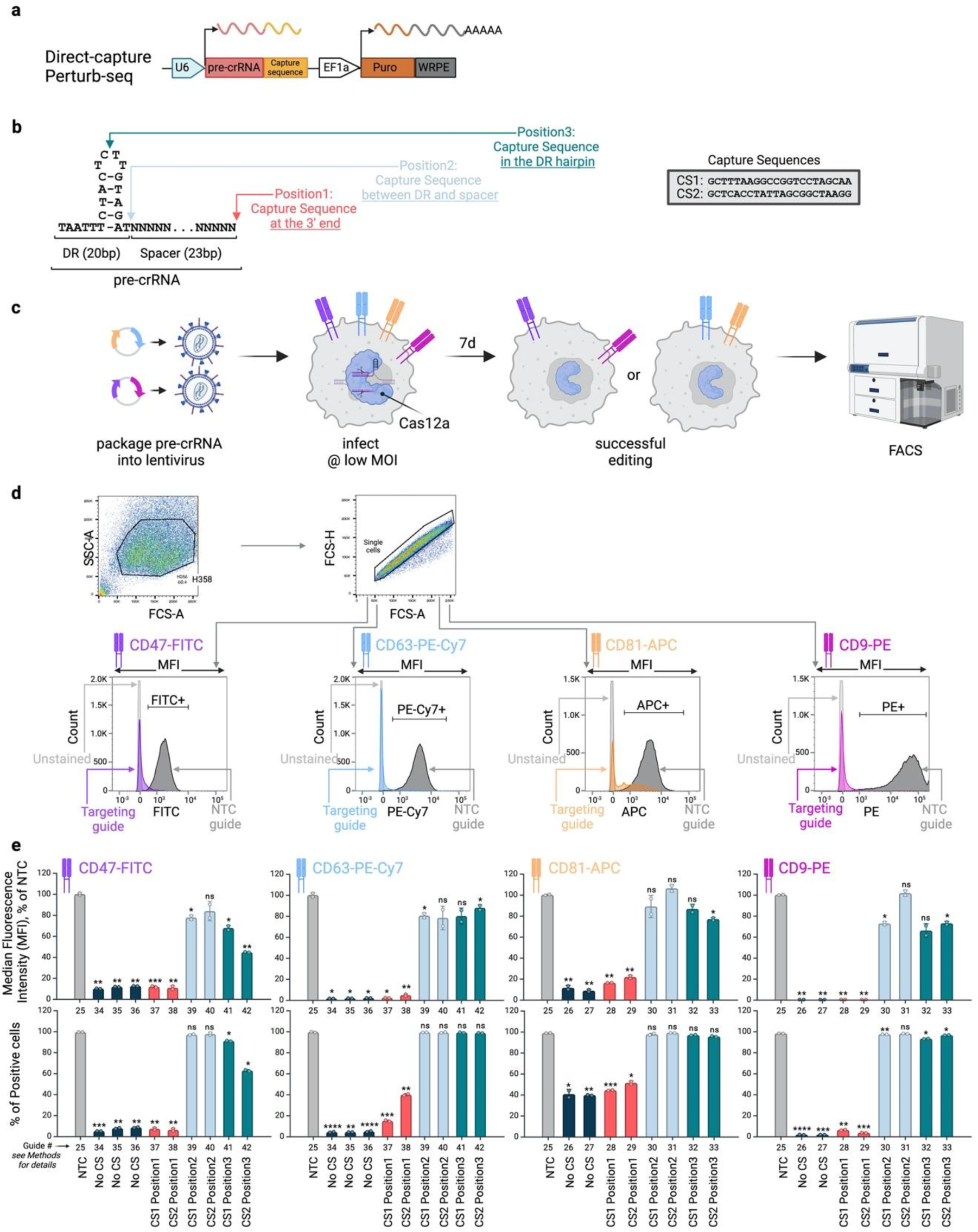
Capture Sequences at Cas12a guide’s 3’ end do not impair guide efficiency. **a,** Schematic of pre-crRNA expressed from direct-capture Perturb-seq formats. **b,** Design of pre-crRNA with capture sequences added at three different positions. **c,** Experimental procedure. **d,** Example of FACS cell gating strategy to record mean fluorescence intensity (MFI) and % of positive cells for a particular channel. **e,** Quantification of FACS data. Only capture sequences (CS) at the 3’ end of the Cas12a guide allows for efficient editing. NTC = non-targeting control. Bar heights represent the Mean of two biological replicates normalized by average NTC values. Error bars represent SD. Statistical comparison between guide formats was performed by an unpaired two-sample t-test with Welch’s correction whereby *p < 0.05, **p < 0.01, ***p < 0.001, ****p < 0.0001 as compared to the NTC (Guide #25).

**Extended Data Fig. 2.**
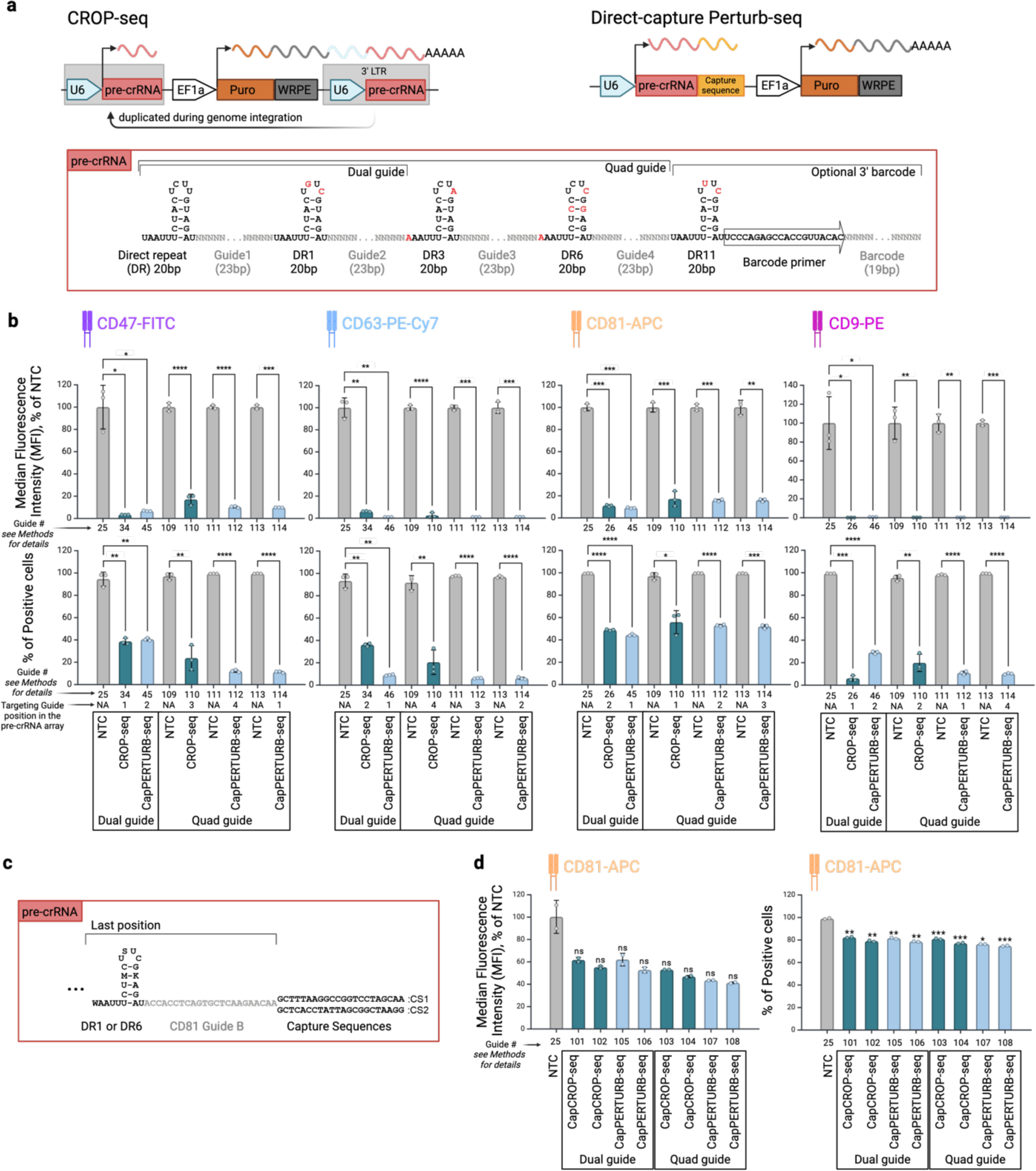
Cas12a performs efficient multiplex targeting with pre-crRNA from different construct formats. **a,** Schematics of pre-crRNA expressing constructs and design of pre-crRNA array with dual and quad guides found within both formats. In CROP-seq pre-crRNA is part of the 3’ LTR, which gets duplicated during lentiviral genome integration. This creates a second copy of pre-crRNA, which produces functional Pol III transcripts. pre-crRNA that is part of long Pol II puro^R^ transcript is the version detected in scRNA-seq from its polyA. The U6 promoter downstream of puro^R^ has been shown not to produce RNA. However, unlike Cas9, Cas12a can probably process crRNAs out of the long Pol II transcript, which would increase the amount of available guide RNA. **b,** Quantification of FACS data obtained from comparing dual and quad pre-crRNA arrays in CROP-seq and direct-capture Perturb-seq (CapPERTURB-seq) formats. NTC = non-targeting control. Bar heights represent the Mean of three biological replicates normalized by average NTC values. Error bars represent SD. Statistical comparison between guide formats was performed by an unpaired two-sample t-test with Welch’s correction whereby *p < 0.05, **p < 0.01, ***p < 0.001, ****p < 0.0001 as compared to the NTC (Guides #25, 109, 111, 113). **c,** Design of the last position in a pre-crRNA array expressing a sub-optimal CD81 guide followed by a capture sequence. **d,** Quantification of FACS data obtained with the pre-crRNA set-up from panel c. Bar heights represent the Mean of two biological replicates normalized by average NTC values. Error bars represent SD. Statistical comparison between guide formats was performed by an unpaired two-sample t-test with Welch’s correction whereby *p < 0.05, **p < 0.01, ***p < 0.001, ****p < 0.0001 as compared to the NTC (Guide #25).

**Extended Data Fig. 3.**
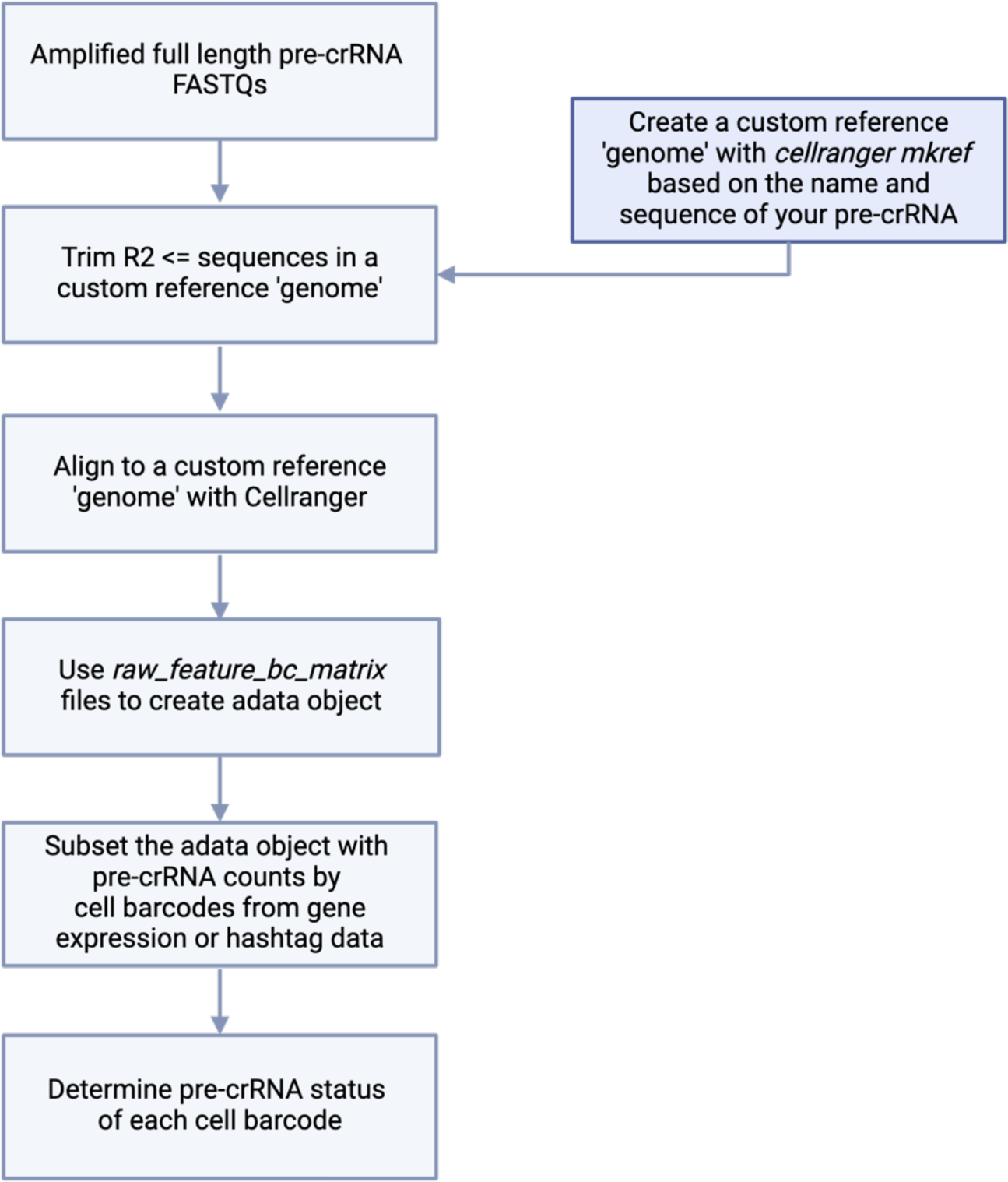
Bioinformatic workflow for identification of pre-crRNA. Schematic of workflow used to identify pre-crRNA whereby reads (R2) from FASTQ files from a library of amplified full-length pre-crRNA are trimmed (if necessary) and aligned to a custom reference genome using 10x Genomics Cell Ranger v7.1.0. Custom reference genome is also generated by Cell Ranger based on pre-crRNA sequences in the library with mkref function. Post reads alignment, the raw feature barcode matrix file is filtered to extract pre-crRNA counts by cell barcode allowing for pre-crRNA status assignment to individual cells.

**Extended Data Fig. 4.**
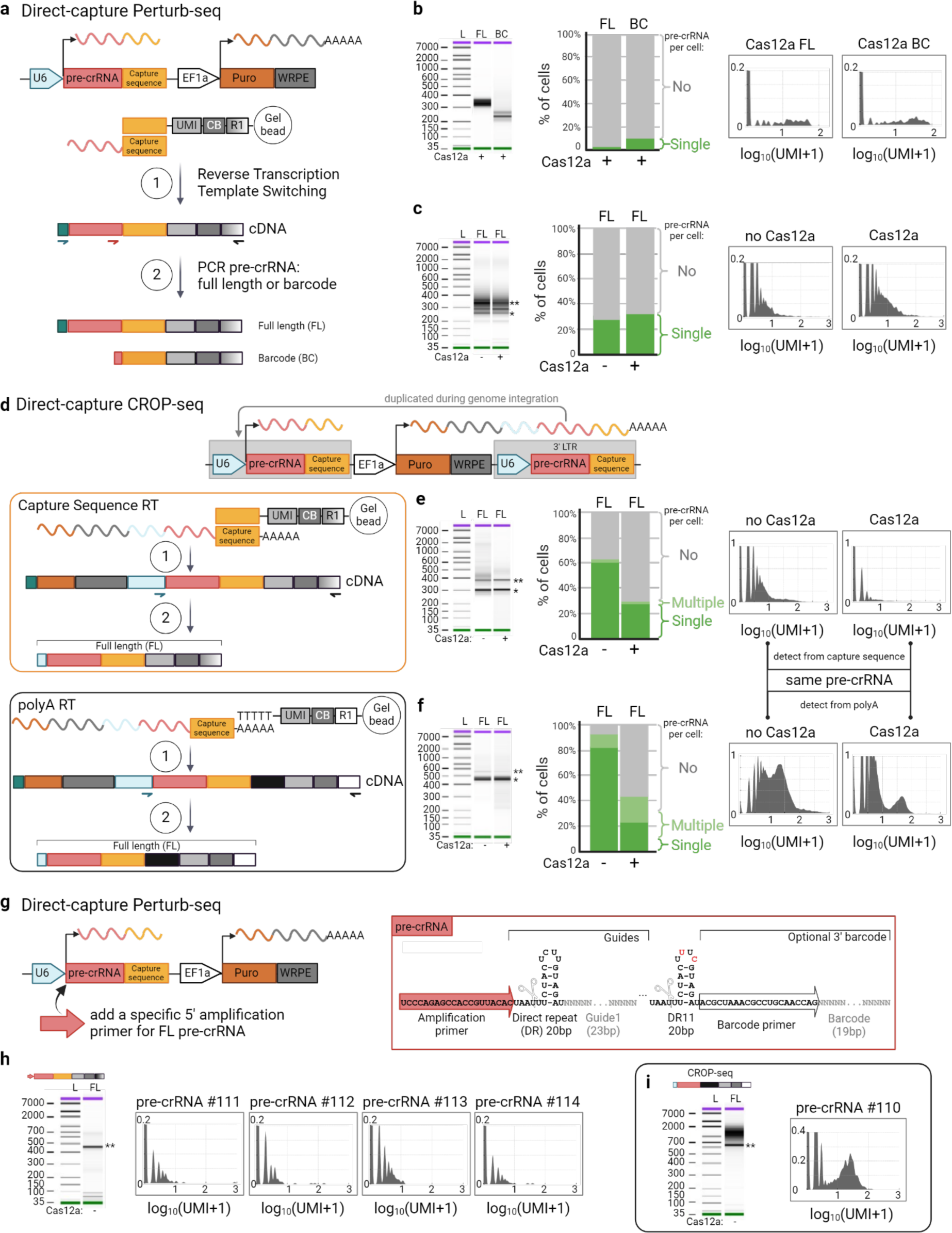
pre-crRNA format optimization. **a,** Scheme showing Direct-capture Perturb-seq construct and protocol for generation of the CRISPR guide libraries, when short capture sequences attached to guide RNA binds to complementary capture sequences part of oligonucleotides attached to magnetic beads. Unlike CROP-seq, direct-capture Perturb-seq constructs do not have a specific 5’ primer landing site and use template switch oligo (TSO) present on all amplified cDNA. Nextera reverse primer can also distinguish direct-capture Perturb-seq from CROP-seq constructs, which are amplified with Illumina reverse primer. **b,** Bioanalyzer traces of epigenetic factor library in direct-capture Perturb-seq format amplified from TSO for full-length (FL) sequences or from primer specific to 3’ end barcode (BC). Percentage of cells identified as expressing none, single, or multiple pre-crRNA constructs. Signal from representative pre-crRNA obtained by full-length and barcode detection methods. **c,** Same as in b but for a set of pre-crRNAs in direct-capture Perturb-seq format targeting the cell-surface receptors. **d,** Construct scheme for direct-capture CROP-seq format with two vignettes differentiating between amplification of the same RNA detected with capture sequences from polyA:polyT interaction. **e,** Bioanalyzer traces, percentage of cells and signal from representative pre-crRNA obtained from direct-capture CROP-seq constructs amplified from capture-sequences as shown in the opposite scheme in d. **f,** Same as in e but for the amplification from polyA:polyT interaction. **g,** Addition of specific forward amplification primer to the 5’ of pre-crRNA in direct-capture Perturb-seq format. **h,** Bioanalyzer trace and signal from pre-crRNAs in the format outlined in g. **i,** Bioanalyzer trace and signal from pre-crRNAs in CROP-seq format run alongside the pre-crRNAs shown in h. L = ladder. * marks a band for dual-guide constructs, ** - quad-guide constructs.

**Extended Data Fig. 5.**
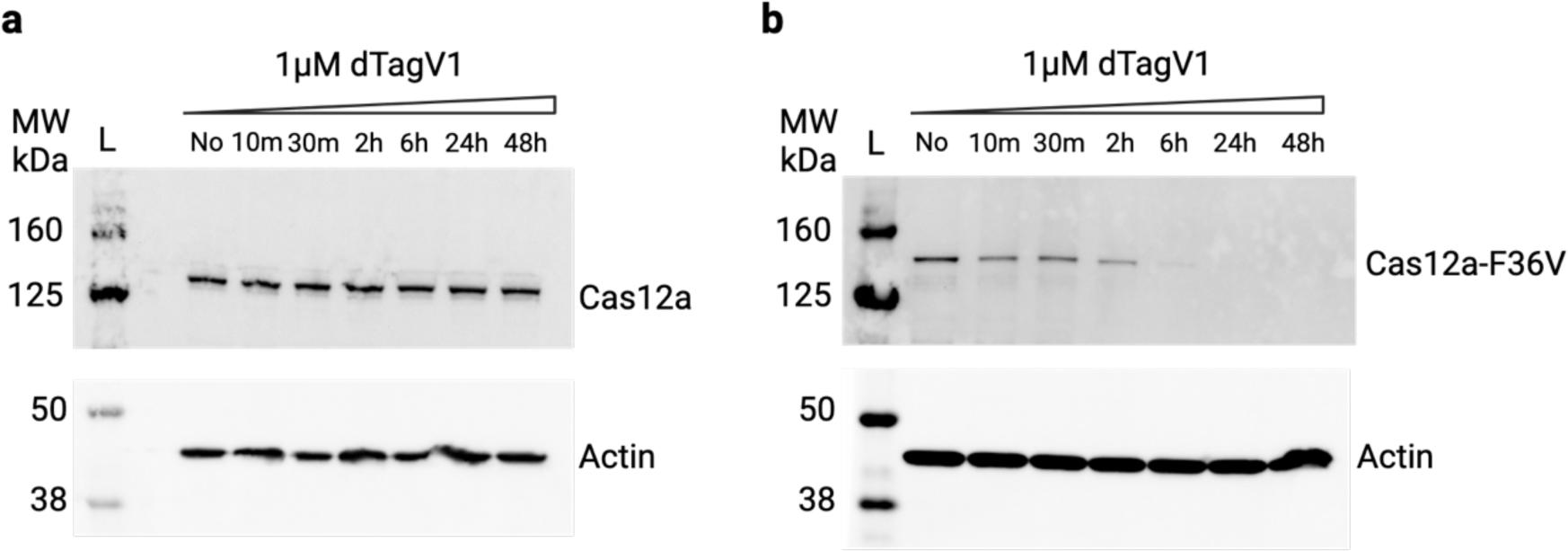
Western blot of Cas12a levels upon dTagV1 addition to induce its degradation. **a,** H358 cells expressing enAsCas12a without FKBP12^F36V^ domain show no degradation of Cas12a with 1µM dTagV1 treatment. Sample labeled ‘No’ was treated with DMSO. **b,** H358 cells expressing enAsCas12a fused to FKBP12^F36V^ (Cas12a-F36V) show progressive degradation of Cas12a with 1µM dTagV1 treatment. Cas12a disappears to levels below detection after 6 hours incubation with the chemical. Sample labeled ‘No’ was treated with DMSO.

**Extended Data Fig. 6.**
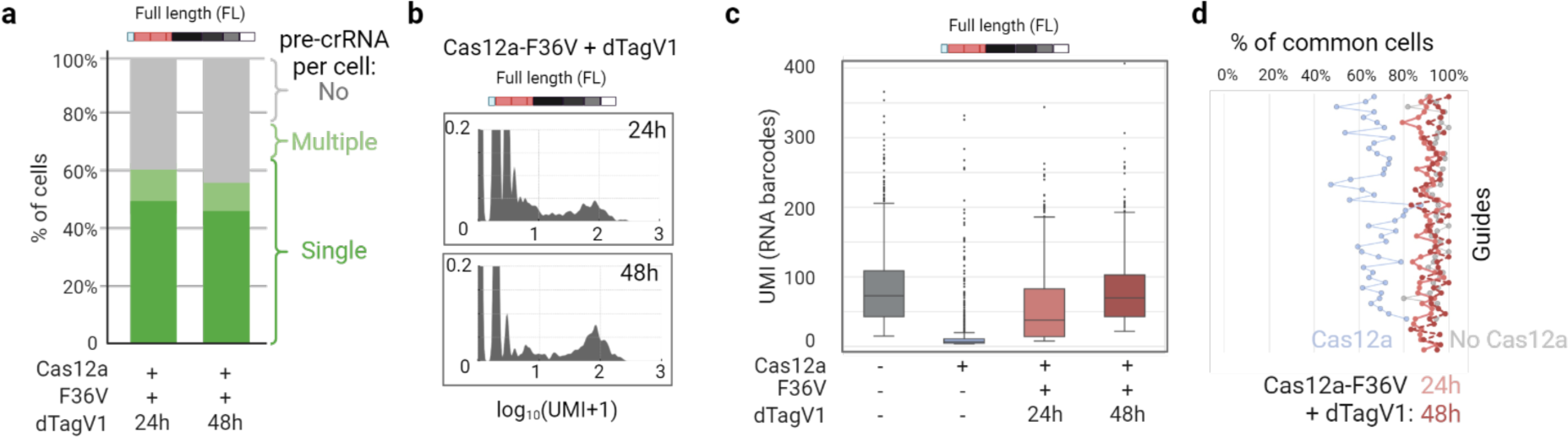
Full length pre-crRNA counts. **a,** Percentage of cells identified as expressing none, single, or multiple pre-crRNA constructs from full-length detection method. **b,** Signal from representative pre-crRNA. **c,** Boxplot of UMI signals for all pre-crRNA in the library. Statistical comparison between samples was performed by an Mann-Whitney U test whereby the p-values were 0 (Cas12a), 1.57e^-78^ (24h), and 0.9193 (48h), as compared to the first sample (no Cas12a). **d,** Percentage of cells with consistent pre-crRNA identify calls between full-length and barcode methods after 24h or 48h dTagV1 treatment overlaid over previously shown data from cells with and without Cas12a. Degradation of Cas12a restores consistency in cell identity calls to the levels of cells without Cas12a.

**Extended Data Fig. 7.**
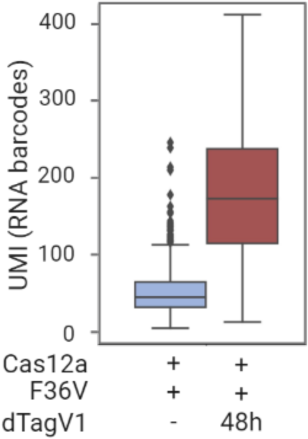
pre-crRNA counts in K562 cells. Boxplot of UMI signals for pre-crRNAs obtained from CROP-seq vectors expressed in K562 cells. Similar to H358 cells, degradation of Cas12a recovers pre-crRNA counts. Statistical comparison between samples was performed by an Mann-Whitney U test whereby the p-value was 1.2e^-278^.

**Extended Data Fig. 8.**
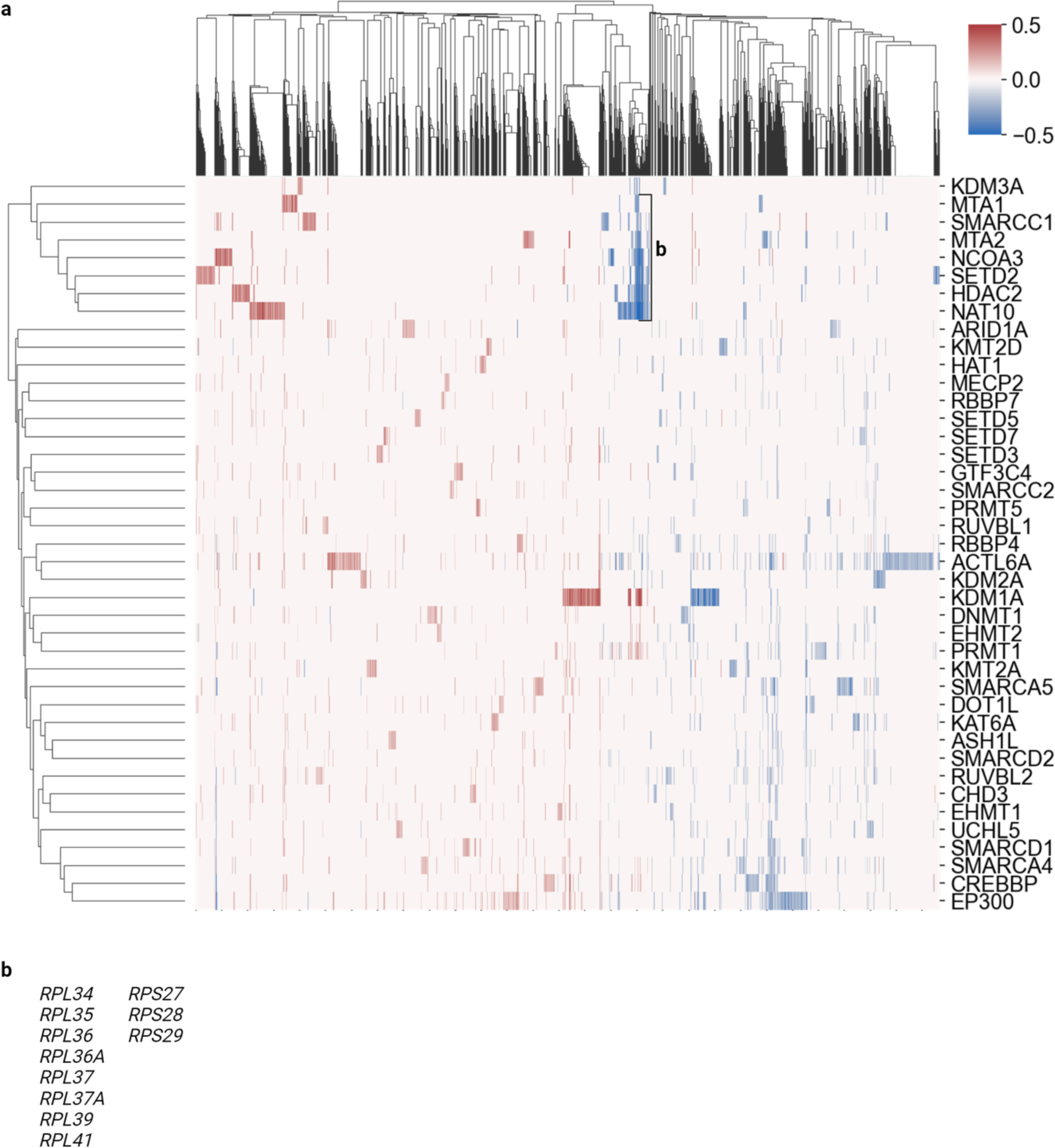
Epigenetic factor screen details. **a,** Heatmap of gene expression changes in response to genetic perturbations from Perturb-seq data. Values correspond to log fold changes in expression of profiled genes (columns, g=12,522) resulting from various genetic perturbations (rows). The dendrogram on the x-axis clusters perturbations with similar gene expression profiles, while the dendrogram at the top clusters genes with similar response patterns across perturbations. *KDM1A* and *NAT10* perturbations are highlighted in the main text with the additional examples following here. **b,** Ribosomal proteins commonly downregulated among several perturbations as highlighted on the heatmap in a including cells with *NAT10* pre-crRNA featured in Fig. 2.

### Supplemental Note 1

There were no observable differences in targeting efficacy when comparing the CROP-seq and direct capture Perturb-seq guide RNA vectors expressing either two or four gene-specific crRNAs per vector (Extended Data Fig 2b). However, we reasoned that the dynamic range of those assays was limited due to the high efficiency of our selected guides. Also, while others have suggested there are minimal position effects for pre-crRNA arrays with ≤4 co-expressed guides, we could not exclude that possibility with the single cell-formatted vectors through the use of these particular spacers.

To further assess differences in efficiency between vector formats and position within a pre-crRNA array, we designed a set of pre-crRNAs with the last position occupied by a new CD81 spacer that induced relatively-low target depletion (Extended Data Fig. 2c). We reasoned that a spacer with ∼50% efficiency would allow us to better discriminate between crRNA expression contexts. We also added capture sequences to the 3’ end of pre-crRNA in the CROP-seq vector, which created the direct-capture CROP-seq format.

We measured CD81 knock-out efficiency with constructs expressing the less efficient spacer and again detected similar targeting efficiency between dual and quad pre-crRNA variants in both the CROP-seq and Perturb-seq formats (Extended Data Fig. 2d).

Together, these data show that CROP-seq and direct-capture Perturb-seq formats exhibit comparable Cas12a-based target disruption with no significant differences between dual and quad pre-crRNA formats. In the direct-capture Perturb-seq format, we found that capture sequences (CS) at the 3’ end of the spacer performed best while other positions interfered with targeting. Additionally, we found that knock-out efficiency was more influenced by the spacer sequences than by the pre-crRNA formats.

### Supplemental Note 2

scRNA-seq of pre-crRNAs expressed from the direct-capture Perturb-seq format resulted in poor guide recovery, whether or not the cells contained enAsCas12 and with different batches of encapsulating beads from 10X (Extended Data Fig. 4a-c).

We hypothesized that the pre-crRNA length in the direct-capture Perturb-seq format was too short for proper detection. To address this, we adopted a direct-capture CROP-seq format, where capture sequences were appended to the pre-crRNA cassette in CROP-seq constructs. This permitted guide RNA transcript complementarity to oligo(d)T or cognate capture sequences and amplification with either of two different reverse primers (Illumina vs Nextera sequences) during scRNA-seq library preparation. This separated RNAs bound through polyA:polyT interactions or from the capture sequences (see schema in Extended Data Fig. 4d). Upon comparing the two RNA capture methods for the same constructs, we observed that sample preparation using the oligo(d)T-based approach was superior to the use of specific capture sequences (Extended Data Fig. 4e, f).

We noticed that PCR amplification of full length pre-crRNAs from direct-capture Perturb-seq constructs did not produce a discrete product of the expected size (Extended Data Fig. 4b,c) and reasoned that this could be due to the template switch oligo (TSO) used as one of the primers, thereby leading to off-target amplification and reduced pre-crRNA detection. To address this, we added a unique primer binding sequence to the 5’ end of the pre-crRNA upstream of the first direct repeat. Upon expression in context with the guide, this sequence would be cleaved off by enAsCas12a during crRNA processing (Extended Data Fig. 4g).

Priming from this sequence resulted in a specific amplicon for the full-length pre-crRNA (Extended Data Fig. 4h). However, the improved full-length pre-crRNA amplification quality did not enhance detection from direct-capture Perturb-seq format (Extended Data Fig. 4h), which was again poor compared to the CROP-seq format used in a parallel scRNA-seq experiment (Extended Data Fig. 4i).

Although we achieved comparable results in targeting efficiency for multiple cell surface receptors in bulk for both CROP-seq and direct-capture Perturb-seq formats, the CROP-seq variant offers a streamlined approach through standard oligo(d)T-based enrichment with efficient and reliable detection of pre-crRNAs at the single-cell level.

